# Deep Learning Based BCI Control of a Robotic Service Assistant Using Intelligent Goal Formulation

**DOI:** 10.1101/282848

**Authors:** D. Kuhner, L.D.J. Fiederer, J. Aldinger, F. Burget, M. Völker, R.T. Schirrmeister, C. Do, J. Bödecker, B. Nebel, T. Ball, W. Burgard

## Abstract

As autonomous service robots become more affordable and thus available for the general public, there is a growing need for user-friendly interfaces to control these systems. Control interfaces typically get more complicated with increasing complexity of the robotic tasks and the environment. Traditional control modalities as touch, speech or gesture commands are not necessarily suited for all users. While non-expert users can make the effort to familiarize themselves with a robotic system, paralyzed users may not be capable of controlling such systems even though they need robotic assistance most. In this paper, we present a novel framework, that allows these users to interact with a robotic service assistant in a closed-loop fashion, using only thoughts. The system is composed of several interacting components: non-invasive neuronal signal recording and co-adaptive deep learning which form the brain-computer interface (BCI), high-level task planning based on referring expressions, navigation and manipulation planning as well as environmental perception. We extensively evaluate the BCI in various tasks, determine the performance of the goal formulation user interface and investigate its intuitiveness in a user study. Furthermore, we demonstrate the applicability and robustness of the system in real world scenarios, considering fetch-and-carry tasks and tasks involving human-robot interaction. As our results show, the system is capable of adapting to frequent changes in the environment and reliably accomplishes given tasks within a reasonable amount of time. Combined with high-level planning using referring expressions and autonomous robotic systems, interesting new perspectives open up for non-invasive BCI-based human-robot interactions.

## 1. Highlights

- BCI-controlled autonomous robotic service assistant
- First online brain-computer-interface using deep learning
- Menu-driven language generation based on referring expression
- Modular ROS-based mobile robot interaction
- Experimental evaluation using a real robot

## 2. Introduction

Persons with impaired communication capabilities, such as severely paralyzed patients, rely on constant help of human care-takers. Robotic service assistants can re-establish some degree of autonomy for these patients, if they offer adequate interfaces and possess a sufficient level of intelligence. Generally, such systems require adaptive task- and motion-planning modules to determine appropriate task plans and motion trajectories for the robot to execute a task in the real world. Moreover, it requires a perception component to detect objects of interest or to avoid accidental collisions with obstacles. With increasing capabilities of autonomous systems intelligent control opportunities also become more important. Typical interfaces, such as haptic (buttons), audio (speech) or visual (gesture) interfaces, are well suited for healthy users. However, for persons with impaired communication skills these control opportunities are unreliable or impossible to use.

In this paper, we present and evaluate a novel framework, schematically depicted in Fig. 1, that allows closedloop interaction between users with minimal communication capabilities and a robotic service assistant. To do so, we record neuronal activity elicited in the human brain, the common origin of all types of communication, with an electroencephalography (EEG) system. Furthermore, we employ a deep convolutional neural network (ConvNet) approach for online co-adaptive decoding of neuronal activity, in order to allow users to navigate through a graphical user interface (GUI) which is connected to a high-level task planner. It allows the intuitive selection of goals based on the generation of referring expressions that identify the objects to be manipulated. The set of feasible actions displayed in the GUI, depends in turn on the current state of the world, which is stored in a central knowledge base and continuously updated with information provided by the robot and a camera perception system. Once a task has been selected, it is decomposed into a sequence of atomic actions by the high-level planner. Subsequently, each action is resolved to a motion for the mobile manipulator using low-level motion-planning techniques. This approach minimizes the cognitive load required of the user, which is a crucial aspect in the design of a BCI. Furthermore, the intelligence and autonomy of the system make it possible to interface non-invasive BCIs, which currently have low throughput, with our robotic assistant composed of 11 degrees-of-freedom (DOF). In the following, we present the related work, describe the individual components shown in Fig. 1 and present a quantitative evaluation of the system regarding its performance and user-friendliness.

**Figure 1:**
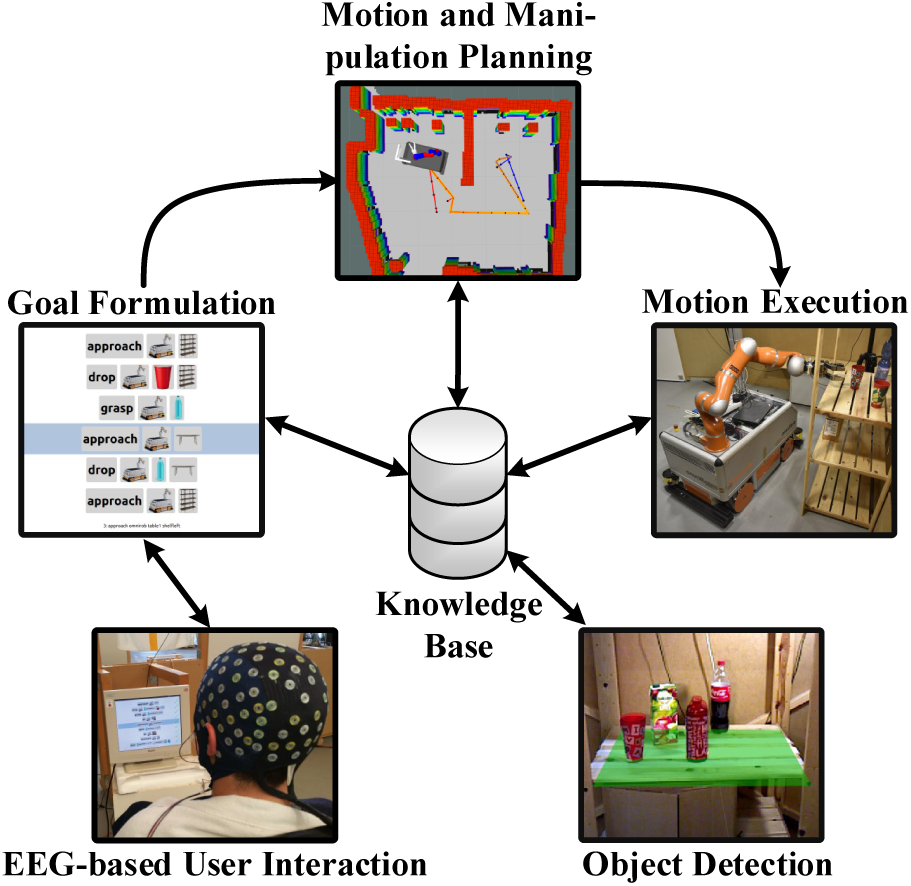
Our framework that unifies decoding of neuronal signals, high-level task planning based on referring expressions, low-level motion- and manipulation-planning, and scene perception with a centralized knowledge base at its core. Intuitive goal selection is provided through an adaptive graphical user interface.

## 3. Related Work

The multi-disciplinary work presented in this paper relies on robotics, brain-computer interfaces and natural-language generation (NLG). This section outlines related work in these fields.

### Robotic Assistants

Multiple previous studies have focused on robotic systems assisting people with disabilities. For example, Park *et al*. [1] implemented a system for autonomously feeding yogurt to a person. Chung *et al*. [2] focus on autonomous drinking which involves locating the drink, picking it up and bringing it to the person’s mouth. Using a hybrid BCI and head movement control, Achic *et al*. [3] studies a setup with a moving wheelchair and an attached robotic arm. None of these systems use pure BCI control. In contrast, Wang *et al*. [4] employ a motor imagery BCI with three classes to achieve low-level control of a robotic arm. More relevant, Schröer *et al*. [5] propose a robotic system which receives a BCI command from a user and autonomously assists the user in drinking from a cup. However, this approach only considers a single object and a fixed-base manipulator. Grigorescu *et al*. [6] use steady-state visually evoked potentials to control the commercially available FRIEND III assistance robot. This work is perhaps closest to ours with respect to the number of possible commands (namely 5), the high-level control concept and the (semi-)autonomy of the assistance robot. In contrast to their work, we use active brain signals to control the graphical user interface and apply co-adaptive training and decoding. See the excellent review of Mladenović *et al*. [7] for details on co-adaptive BCIs. Additionally, we propose a specific design of the user interface to improve the human-robot interaction and show that our system is fully autonomous. Most recently, the work of Muelling *et al*. [8] presents a shared-control approach in the field of assistive robotics based on an invasive BCI. This is contrary to our approach which relies on a non-invasive BCI. Nonetheless, their approach could be combined with the goal formulation interface presented in this work.

### Brain-Computer Interfaces

To ensure user acceptance, robust decoding of brain signals is required. Inspired by the successes of deep ConvNets in computer vision [9, 10] and speech recognition [11, 12], deep ConvNets have recently been applied more frequently to EEG brain-signal decoding and related to this paper to decode tasks in brain-computer interfaces. Lawhern *et al*. [13] use a deep ConvNet to decode P300 oddball signals, feedback error-related negativity and two movement-related tasks. In cross-participant evaluation (i. e., trained on some participants and evaluated on others), their ConvNet yields competitive accuracies compared to widely-used traditional brain-signal decoding algorithms. Tabar and Halici [14] combine a ConvNet and a convolutional stacked auto-encoder to decode motor imagery within-participant and improve accuracies compared to several non-ConvNet decoding algorithms. Schirrmeister *et al*. [15] use a shallow and a deep ConvNet to decode both motor imagery and motor execution within-participant. Their approach results in comparable or slightly better accuracies than the widely used EEG motor-decoding algorithm *filter bank common spatial patterns* [16]. Bashivan *et al*. [17] estimate the mental workload with a ConvNet trained on fourier-transformed inputs. In addition to the above work on evaluating ConvNet decoding accuracies, ConvNet visualization methods allow us to get a sense of what brain-signal features the network is using [15, 17, 18, 19, 20, 21]. Taken together, these advances make deep ConvNets a viable alternative for brain-signal decoding in brain-computer interfaces. A first attempt at using shallow ConvNets for online BCI has recently been reported [22]. To the best of our knowledge, apart from our previous paper [23], there is no other work, which uses a deep ConvNet-based online control to implement an EEG-based brain-computer interface.

### Referring Expressions

When humans communicate goals to other humans, they identify objects in the world by referring expressions (e. g., *a red cup on the shelf*). The generation of referring expressions has been subject to computational linguistics research for years as one part of natural language generation (NLG) [24]. With recent advances in natural language processing, computer vision and the rise of neuronal networks, it is nowadays possible to identify objects in images by building referring expressions generated from features [25]. Spatial references can be used to discriminate similar objects [26]. The NLG problem has been approached with planning techniques [27]. However, such systems usually lack knowledge about the actions that can be executed and the objects that can be manipulated. To overcome this problem and to improve the human-robot interaction we propose a user interface that allows specifying actions in a domain-independent way and automatically adapts to changes in the environment.

### Taskand Manipulation-Planning

In contrast to classical task planning, Task- and Manipulation-Planning (TAMP) algorithms also consider the motion capabilities of the robot to determine feasible task plans. There are various approaches to solve the this problem. Common to most TAMP approaches is a hierarchical decomposition of the problem into task- and motion-planning layers. Due to the high dimensionality of the TAMP problem the decomposition can be understood as a way to guide the lowlevel planners based on the high-level plan solution and vice versa. For example, Kaelbling *et al*. [28, 29] propose an aggressively hierarchical planning method. Such a hierarchical decomposition allows handling problems with long horizons efficiently. De Silva *et al*. [30] show an approach based on Hierarchical Task Networks (HTNs) to reason on abstract tasks and combine them with a geometric task planner which works in a discrete space of precomputed grasp, drop and object positions. Recently, the work of Dantam *et al*. [31] introduce the probabilisticallycomplete Iteratively Deepened Taskand Motion-Planning (IDTMP) algorithm, which uses a constrained-based task planner to create tentative task plans and sampling-based motion planners for feasibility tests. Srivastava *et al*. [32] focus on a planner-independent interface layer between task- and motion-planners. Lozano-Pérez *et al*. [33] postpone the decision on motion plans to avoid expensive backtracking due to restrictions which might happen, if the lowlevel planner is queried too early. Instead, they generate a “skeleton” high-level plan and a set of constraints, which need to be satisfied to achieve the goals of the high-level planner. Dornhege *et al*. [34] integrate taskand motionplanning by extending the TFD task planner [35] with semantic attachments, i. e., modules which check the feasibility of motion plans on demand to ensure that task plans can be refined to motion plans. In this work, the goal formulation interface outputs a task plan composed of high-level actions. We assume that these actions can be refined to motions of the robot, if the task plan is consistent with the current world model. Thus, task- and manipulation-planning is also considered in a hierarchical way but we postpone the decision on the actual feasibility of motion plans to reduce the computational effort. Nonetheless, due to the modular structure or our system most of the principles applied in this field could be integrated into our framework as well.

## 4. Autonomous BCI-controlled Service Assistant

In this paper, we present an autonomous robotic service assistant which uses a BCI and an intuitive goal formulation framework to aid users in fetch-and-carry tasks. Our system relies on multiple components which are depicted in Fig. 2. The communication between user and robotic service assistant is established using an EEG-based BCI. It decodes the brain signals using deep ConvNets and is explained in Sec. 4.1. Sec. 4.2 describes the goal formulation assistant that employs referring expressions and a menu-driven user interface to allow an intuitive specification of tasks. These are then processed by a high-level task planner to break them into a set of executable subtasks that are sent to the navigationand manipulationplanning algorithms. Furthermore, we apply a MonteCarlo localization approach to estimate the pose of the robot in the environment. Based on these poses and the map of the environment, the navigation module determines a collision-free trajectory, that allows the robot to move between different locations. Additionally, roadmap-based planning allows to execute manipulation tasks as grasping and dropping objects. More details on motion generation are available in Sec. 4.3. The goal formulation interface and the TAMP algorithms depend on a perception module that dynamically detects relevant objects in the environment. Autonomous drinking capabilities also require that the framework is able to determine the user’s mouth location and the robust estimation of liquid level heights to avoid spilling while serving a drink (Sec. 4.4). Finally, to conduct the data communication a knowledge base connects all components (Sec. 4.5).

**Figure 2:**
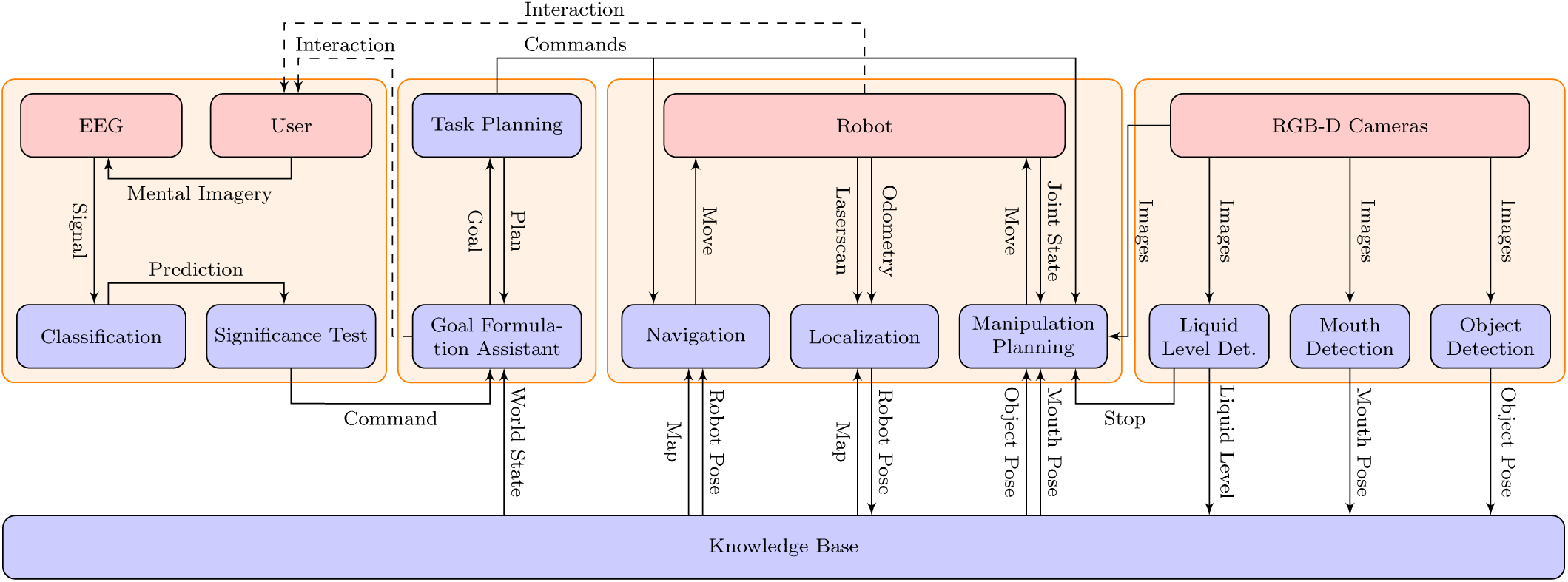
Detailed overview of our framework. It uses a brain-computer interface to decode the thoughts of the user. Thus, the user has control over a goal formulation assistant which is connected to a high-level planner. The commands send by the high-level planner are then processed by the low-level motion planners and executed on the robot. A perception system determines information on object poses, the user’s mouth position and liquid levels. Finally, a central knowledge base stores and provides data to establish a connection between all components.

### 4.1. Online Decoding of Neuronal Signals

This section introduces the deep ConvNet and the stratgies to train the network. Furthermore, we explain the online decoding pipeline to extract meaningful commands from EEG data, which are required to control the robotic assistant.

#### 4.1.1. Deep Hybrid ConvNet Training

As reliable classification of brain signals related to directional commands cannot yet be achieved directly with non-invasive BCIs, we decode multiple surrogate mental tasks from EEG using a deep ConvNet approach [15]. This approach introduces a hybrid network, combining a deep ConvNet with a shallower ConvNet architecture. The deep part consists of four convolution-pooling blocks using exponential linear units (ELU) [36] and max-pooling, whereas the shallow part uses a single convolution-pooling block with squaring non-linearities and mean-pooling. Both parts use a final convolution with ELUs to produce the output features. These features are then concatenated and fed to a final classification layer. All details of the architecture are visualized in Fig. 3.

**Figure 3:**
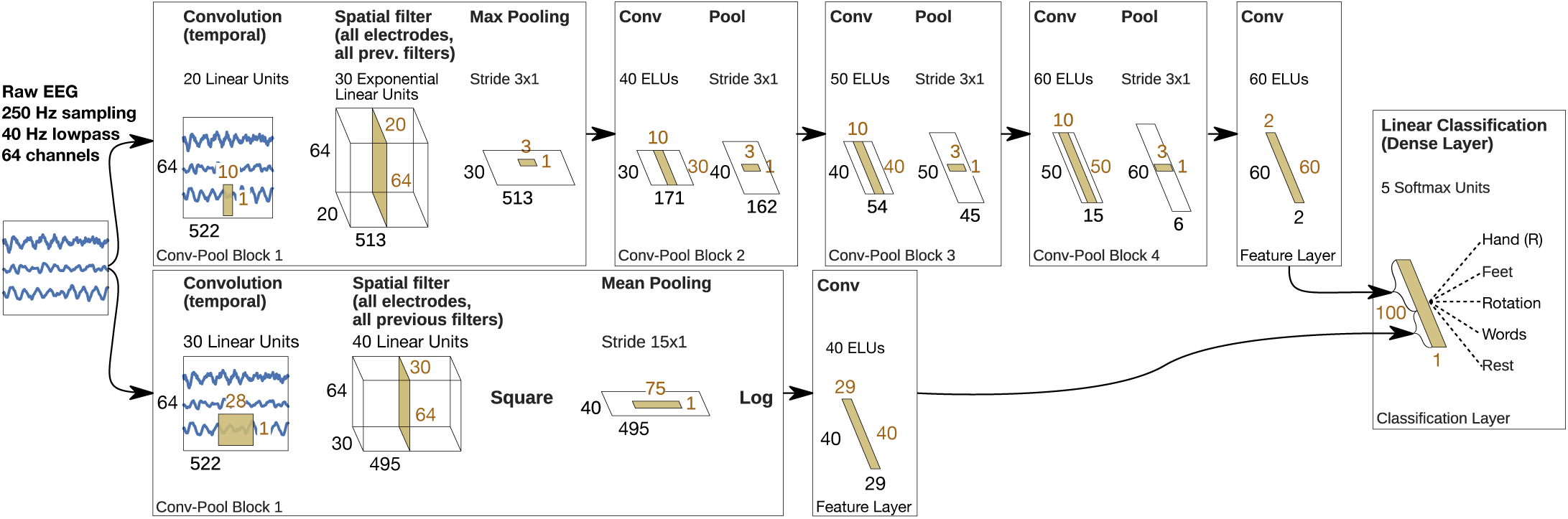
Hybrid deep convolutional neural network: Black numbers depict the dimensions of the input data to each layer. Orange numbers depict the dimensions of the kernels of each layer.

We train the subject-specific ConvNets on 40 Hz lowpass-filtered EEG data to decode five mental tasks: sequential right-hand finger tapping, synchronous movement of all toes, object rotation, word generation and rest. These mental tasks evoke discernible brain patterns and are used as surrogate signals to control the GUI. The mental tasks map to the *select, go down, go back, go up*, and *rest* GUI actions, respectively. *Offline* training is conducted based on a cropped training strategy using shifted time windows, which we call crops, within the trials as input data [15]. The crop size of *∼*2 s (522 samples @ 250 Hz) is given by the size of the ConvNet’s receptive field. Crops start *∼*1.5 s before trial onset and end with trial offset. This corresponds to the first output being predicted 500 ms after trial onset and the last output being predicted on trial offset. To speed up training, one super-crop consisting of 239 consecutive crops (760 samples) is processed in a single forward pass of the model. This results in 239 outputs for each forward pass. One training batch consists of 60 super-crops. We perform stochastic gradient descent using Adam [37] and a learning rate of 10^−3^. To optimize the model we minimize the categorical cross entropy loss. For offline evaluation, we retain the last two runs as our test set, which corresponds to 20 min of data. The remaining data is split into training (80 %) and validation (20 %) sets. We train for 100 epochs in the first phase of the training and select the epoch’s model with the highest validation accuracy. Using the combined training and validation sets, retraining is performed until the validation loss reaches the training loss from the epoch selected in the previous phase. Throughout this paper, we report decoding accuracies on the test set. The initial model used for *online* co-adaptive training is trained on all available *offline* data by following the above mentioned scheme.

We perform *online* co-adaptive training with tenfold reduced learning rate, a super-crop size of 600 samples and a batch size of 45 super-crops. We keep all other parameters identical and train for five batches in all 2 s-breaks. During this time incoming EEG data is accumulated and processed once the newly trained ConvNet is available. Training initiates once ten trials have been accumulated in an experimental session. Only session specific data is used during the training. A session is defined as the time interval during which the participants continuously wear the EEG cap. As soon as the EEG cap is removed and reapplied a new session starts.

#### 4.1.2 Participant Training

Based on our experience it is important to train the BCI decoder and participants in an environment that is as close as possible to the real application environment to avoid pronounced performance drops when transiting from training to application. Therefore, we designed a gradual training paradigm within the goal-formulation user interface (see Sec. 4.2) in which the displayed environment, timing and actions are identical to those of the real control task. The training paradigm proceeds as follows.

##### Offline Training

We first train each participant *offline* using simulated feedback. Participants are aware of not being in control of the GUI. The mental tasks are cued using modified versions of the BCI2000 [38] grayscale images that are presented for 0.5 s in the center of the display. To minimize eye movements the participants were instructed to look at a fixation circle, permanently displayed in the center of the GUI. After a random time interval of 1-7 s the fixation circle is switched to a disk for 0.2 s, which indicates the end of the mental task. At the same time the GUI action (*go up, go down, select, go back, rest*) corresponding to the cued mental task (cf. Sec. 4.1.1) is performed to update the GUI. The *rest* mental task is implicitly taking place for 2 s after every other task^2^. To allow the participant to blink and swallow, every 4th rest lasts 7 s. Fig. 4 gives a graphical overview of the offline training paradigm. To keep training realistic we include a 20 % error rate, i. e., on average every fifth action is purposefully erroneous. We instruct the participants to count the error occurrences to assert their vigilance. This offline data is used to train the individual deep ConvNets as described in Sec. 4.1.1.

**Figure 4:**
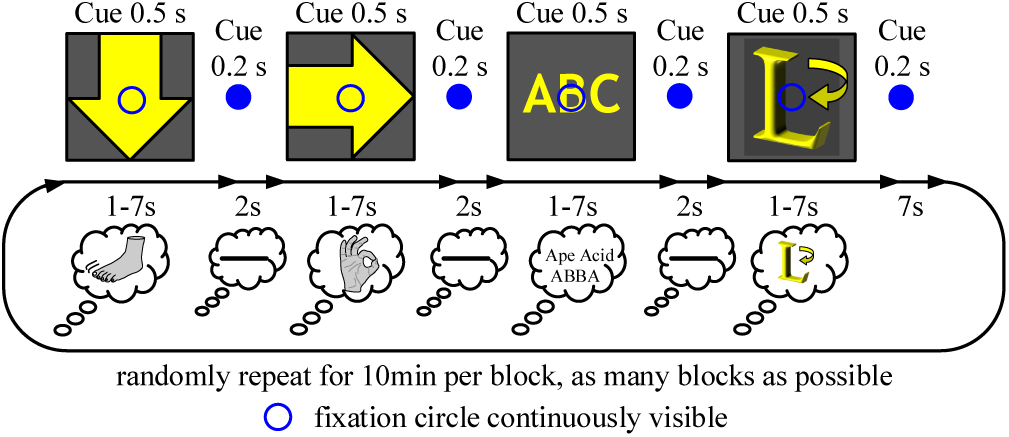
Offline training paradigm. Cue icons, modified from BCI2000 [38], indicate which mental task should be performed by the participant. The cue icons shown here have been modified for better visibility. In our experimental environment we use grayscale versions of the icons. The mental tasks are illustrated by lines in the smaller ‘thought bubbles’. Each mental task maps to a GUI action: word generation *→ go up*, synchronous movement of all toes *→* go down, sequential right-hand finger tapping *→ select*, object rotation *→* go back, rest *→* rest

##### Online Co-Adaptive Training

After offline training, the participants transit to co-adaptive *online* training where the cued mental tasks are decoded by the ConvNets and performed in the GUI. The ConvNets were retrained after each trial during the 2 s break, as described in Sec. 4.1.1. The participants are conscious of being in control of the GUI and are instructed to count the errors they make. In doing so, the participants are aware of their performance, which potentially triggers learning processes and asserts their vigilance.

##### Online Training

To evaluate the *uncued, online* performance of the BCI control, we stop cueing the mental tasks and let the participants select instructed goals in the GUI. The corresponding task plans are then executed by a simulated robot or – when available – the real mobile manipulator. To provide more control over the mobile manipulator and enhance the feeling of agency, participants have to confirm the execution of every planned action and can interrupt the chain of actions at any moment during their execution using the *go back* GUI action. BCI decoding accuracies for the label-less instructed tasks are assessed by manually rating each decoding based on the instructed task steps. Statistical significance of the decoding accuracies are tested using a conventional permutation test with 100 k random permutations of the labels (i. e., the p-value is the fraction of label permutations that would have led to better or equal accuracies than the accuracy of the original labels).

#### 4.1.3 Online Decoding Pipeline

During *online* control of the GUI, the EEG data is lowpass filtered at 40 Hz, downsampled to 250 Hz and sent to a GPU server in blocks of 200 ms for decoding. During coadaptive *online* training (cf. Sec. 4.1.2) the data is additionally labeled (to identify mental tasks) before being sent to the GPU server for decoding, storing and subsequent training. On the GPU server 600 samples (one super-crop, 2.4 s @ 250 Hz) are accumulated until the decoding process is initiated. Subsequently, a decoding step (forward pass of the ConvNet) is performed whenever 125 new samples (0.5 s @ 250 Hz) have accumulated. All predictions are sent back to the EEG-computer on which a growing ring buffer stores up to 14 predictions corresponding to 7 s of EEG data. Once the ring buffer contains two predictions (i. e., 1 s) our algorithm extracts the mental task with the largest mean prediction. A two-sample t-test is then used to determine if the predictions significantly deviate from 0.05. We define significance as *p <* 0.2^3^. These two steps are repeated for all predictions until significance is reached. The ring buffer’s size increases (max. 14 predictions) as long as the predictions are not significant. Once significance is reached the GUI action linked to the mental task is executed and the ring buffer is cleared.

### 4.2. Goal Formulation Planning

Our approach adopts domain-independent planning for high-level control of the robotic system. Whereas many automated planning approaches seek to find a sequence of actions to accomplish a predefined task, the intended goal in this paper is determined by the user. Specifying goals in the former case requires insight into the internal representation of objects in the planning domain. By using a dynamic knowledge base that contains the current world state and referring expressions that describe objects based on their type and attributes, we obstruct direct user access to the internal object representation. Furthermore, we are able to adapt the set of possible goals to changes in the environment. For this purpose, our automatic goal formulation assistant incrementally builds references to feasible goals in a menu-driven graphical user interface.

#### 4.2.1 Domain-Independent Planning

Automated planning is used to transfer a system into a desired goal state by sequentially executing high-level actions. A planning task consists of a planning domain *𝒟;* and a problem description Π. The former is a tuple *D* = *(𝒯;*, *𝒞;*_*d*_, *𝒫;, 𝒪)*, where

- *𝒟=(𝒯,𝒞*_*d*_,*𝒫,𝒪)*, is the type system together with a partial ordering that specifies the sub-type relations between types in *T*,
- *𝒞*_*d*_ contains a set of domain constant symbols,
- *𝒫* is the set of predicate symbols, and
- *𝒪*corresponds to the set of planning operators and specifies their effects and preconditions.

The problem description Π =*(𝒟; 𝒞t 𝒮)*is defined as fol-lows:

- *𝒟;* is the domain description,
- *𝒞*_*t*_ are the additional task-dependent constant symbols, where *𝒞*_*d*_ *∩ 𝒞*_*t*_ = *∅*, and
- *𝒮* is the initial state.

We specify *𝒟;* and Π using the Planning Domain Definition Language (PDDL) [39]. For example, in the service assistance domains that we use in our experiments, *𝒯;*T contains a type hierarchy, where *furniture* and *robot* are of super-type *base*, and *bottle* and *cup* are of super-type *vessel*. Furthermore, P specifies attributes or relations between objects, e. g., *arm*–*empty* is an attribute indicating whether the robot’s gripper is empty and *position* is a relation between objects of type *vessel* and *base*. Finally, defines actions as *grasp* and *move*. The problem description specifies the initial state *𝒪* including object instances, such as *cup01* of type *cup* and *shelf02* of type *shelf* as well as relations between them, e. g., the *position* of *cup01* is *shelf02*, as illustrated in Fig. 5.

**Figure 5:**
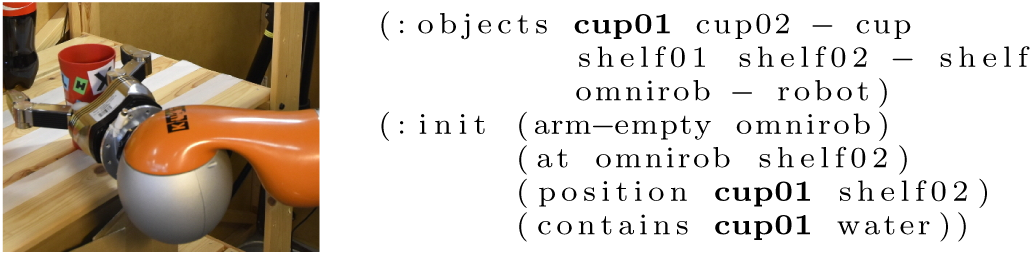
*Left* : The red cup in the real world, referred to by *cup01. Right* : Exemplary PDDL problem description with objects and their initial state.

#### 4.2.2 Human and Machine Understandable References

A major challenge when trying to communicate goals to the user is the limited shared vocabulary between the user and the planning system, whose world is described by a PDDL planning task. The planner’s most concise representation of the cup in Fig. 5 might be *cup01*, which is not sufficiently clear for the user if there are multiple cups. To solve this problem, the goal generation and selection component uses a set of basic references shared between planner and user. These *shared references* can be combined to create *referring expressions* to objects or sets of objects in the world [40, 41]. Generally, a referring expression is a logical formula with a single free variable. We say that *refers* to an object *o* if = (*o*), i. e., is valid in our PDDL domain theory. For example, we can refer to *cup01* by (*x*) *cup*(*x*) *contains*(*x, water*). We restrict ourselves to references that are conjunctions of relations *R*_0_, *…, R*_*m*_ and only allow existential quantifiers, i. e.,

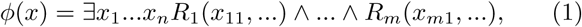

where each argument *x*_*ij*_ corresponds to one of the variables *x*_1_, *…, x*_*n*_. This is preferable for computational reasons and also allows us to incrementally refine references by adding constraints, e. g., adding *contains*(*x, water*) to *cup*(*x*) restricts the set of all cups to the set of cups containing water. A reference *03C6*(*o*) to an object *o* is *unique* if it refers to exactly one object:

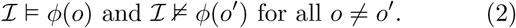

However, it is usually sufficient to create ferences to 432 sets of objects, e. g., if the user wants a glass of ater it 433 might not be necessary to refer to a specific glass as ng 434 as it contains water.

To reference objects in planning domains, we need to specify the components that are required to create *shared references*. We distinguish three fundamental reference types. **Individual references** describe objects that can be identified by their name, e. g., the *content* objects *water* or *apple*-*juice*, and the *omniRob* robot. Additionally, **type name references** are used to specify objects by their type. They allow referring to unspecific objects as a *shelf* or a *cup*. With **relational references** we can refer an object using a predicate in which the object occurs as an argument. In our scenario, most relational references are binary attribute relations whose first parameter is the object that is referred to, and the second parameter is an object in the domain of attribute values. In the example above, a cup can be described using its *content* by the binary relation *contains*(*x, water*).

The most natural way for the planner to represent a goal is a conjunction of predicates, e. g., *cup*(*x*) *shelf* (*y*) *position*(*x, y*) to put a cup on a shelf. This, however, is a rather unnatural way to refer to goals for humans. We found that it is more natural to use the action that achieves the goal than the goal itself, e. g., *action*(*put*, *x, y*) *cup*(*x*) *shelf* (*y*). Therefore, we include **action references**, a macro reference for all predicates in the action’s effect, as additional building blocks to create references to objects in the world and allow the users to specify their goals.

#### 4.2.3 Adaptive Graphical Goal Formulation Interface

In our aim for a flexible yet user-friendly control method to set the robot’s goals, we use the references presented in Sec. 4.2.2 to create a dynamic, menu-driven goal formulation user interface. We allow the user to incrementally refine references to the objects which occur as parameters of a desired goal. We distinguish three different levels of atomicity for the control signals of the GUI: a *step* is a directional command (i. e., *go up, go down, select, go back*) whereas a *stride* is the selection of one refine ment option offered by the GUI. A *stride* is therefore a sequence of *steps* ending with either *go back* (to go back to the previous refinement level) or *select* (to further refine the reference to the object) which does not account for the *go up* and *go down* steps. Finally, a *parameter refinement* is the creation of a reference to one parameter. The goal selection procedure is depicted in Fig. 6. After the initial selection of a goal type, e. g., *drop* (Fig. 6.a), we have to determine objects for all parameters of the selected goal. We start by populating the action with the most specific reference that still matches all possible arguments, e. g., *omniRob, transportable*(*x*) and *base*(*y*), assuming that *omniRob* corresponds to an individual reference and *transportable* and *base* are type-name references (Fig. 6.b). The current goal reference is displayed in the top row of the GUI. The user interface then provides choices to the user for further refinement of the argument. In our example, the first argument *omniRob* is the only object in the world that fits the parameter type *robot* which is why it does not have to be refined any further. Therefore, we start by offering choices for refining the second argument *transportable*(*x*) which yields the selections *bottle*(*x*), *glass*(*x*), *cup*(*x*) and *vase*(*x*). This continues until the argument is either unique, it is impossible to further constrain the argument or any remaining option is acceptable for the user. In the example, we refine the first choice *bottle*(*x*) based on its *content* (Fig. 6.c) by adding a relation *contains*(*x*, *o*) to the referring expression, where *o* is an object of type *content*. This procedure is repeated for all parameters of the goal, which will finally result in a single goal or set of goals (if the references are not unique) that are sent to the high-level planner.

**Figure 6:**
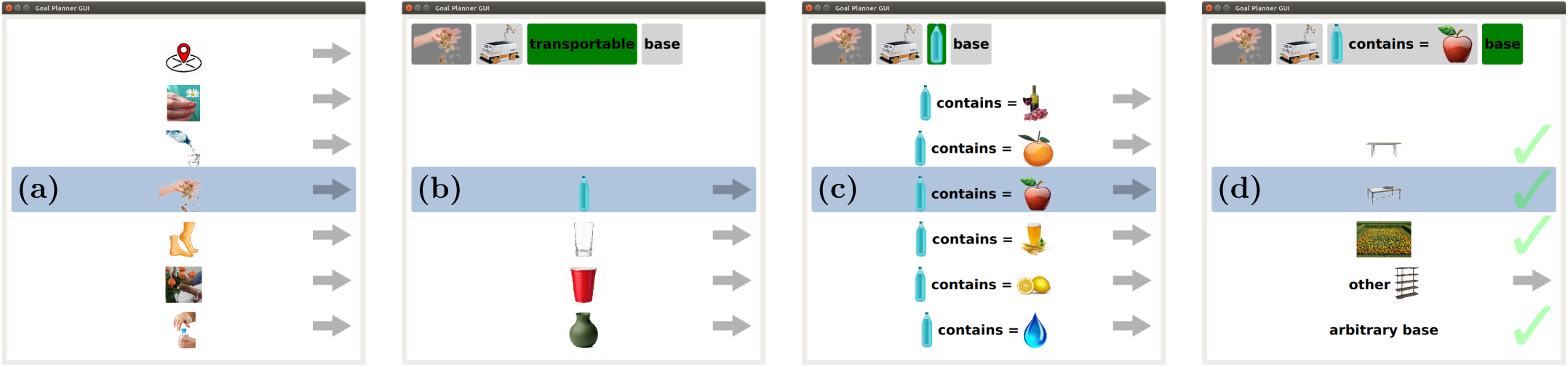
Graphical user interface of the goal formulation assistant. **(a)** Selection of the desired action. **(b)** Refinement of the first action parameter of type *transportable*. **(c)** Refinement of the argument based on *content*. **(d)** Refinement of the last action parameter of type *base*

**Figure 7:**
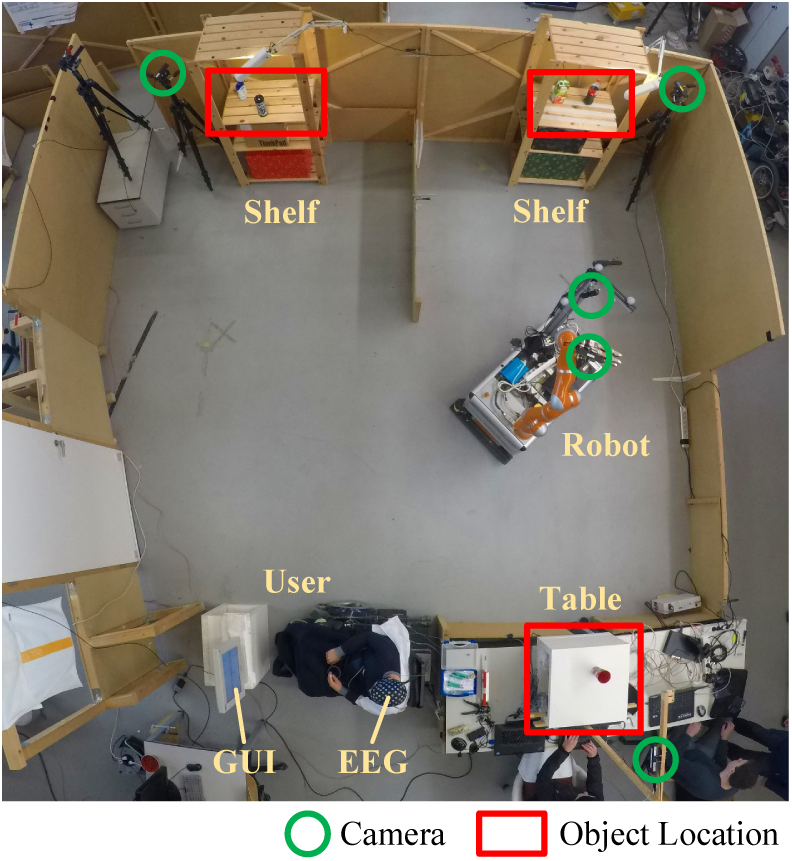
Physical experimental environment: Two shelves and a table could be considered by the robot for performing manipulation actions. Five RGBD sensors observed the environment. A human operator selected a goal using EEG control and the high-level planner GUI.

**Figure 8:**
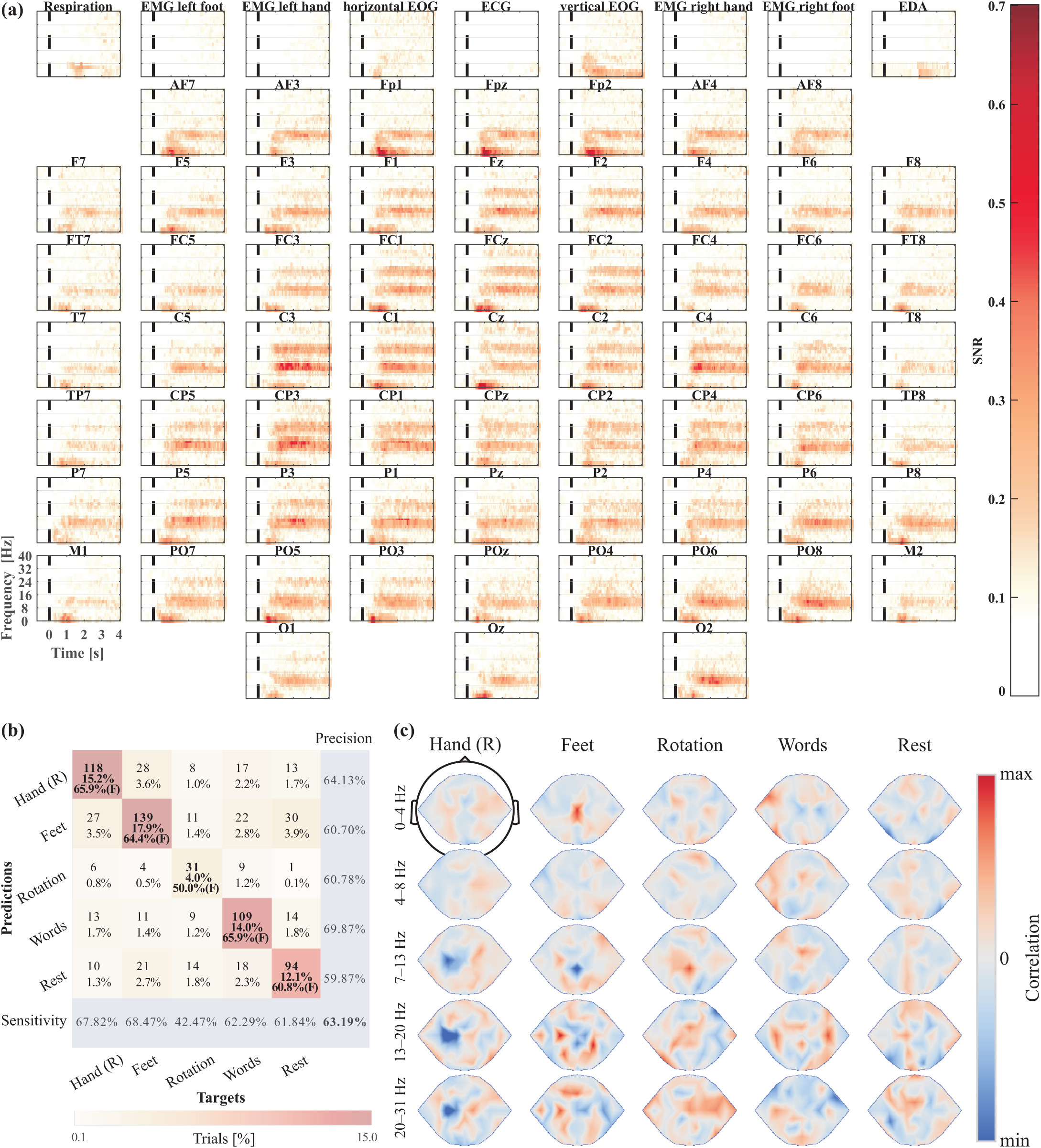
Offline EEG data, offline decoding results and learned features. **(a)** Participant-averaged SNR of the first 4 s of data used to train the hybrid ConvNet. The dashed line indicates the time at which the participants were instructed to start a mental task. Highest SNR can be observed in the alpha (7-14 Hz) and lower beta (16-26 Hz) bands. These frequency bands are robust markers of task related mental activity. Note that the non-EEG channels (top row) were withheld from the ConvNets at any time and are displayed as negative control. The position of most channels was adjusted to achieve a compact layout. **(b)** Confusion matrix of decoding accuracies for the offline train/test transfer pooled over all subjects. Numbers indicate the amount of trials in each cell. Percentages indicate the amount of trials in each cell relative to the total number of trials. (F) indicates F1 score. Dark/light colors indicate that a large/small portion of the targets were predicted for a given class, respectively. **(c)** Topographically plausible input-perturbation network-prediction correlation maps in the delta (0-4 Hz), theta (4-8 Hz), alpha (7-13 Hz), low beta (13-20 Hz) and high beta (20-31 Hz) frequency bands averaged over all participants. The colormap is scaled individually for every frequency band. For details on the visualization technique we refer the reader to [15].

Some features cannot be used to partition the remaining objects for one parameter (e. g., not all objects have the attribute *color*), in which case an entry for all *other* objects can be chosen. Additionally, we allow to skip the refinement of the current parameter and use an *arbitrary* object for it. Finally, we provide an entry to go *back* to the previous refinement stride.

In each stride the shown refined references form a partition, which corresponds to a set of references *ϕ*_*i*_:

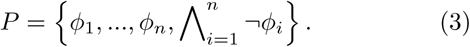

The last term ensures, that the partition covers all objects of the previous reference.

The most important part of the refinement process is to compute possible successors that split the current partition. To make progress in selecting a goal, the successor reference needs to be strictly narrower than its parent. Additionally, forming a partition requires that the references in a candidate set are disjoint. The decision if a complete partition exists corresponds to the NP-complete ExactCover problem. However, by applying a greedy search algorithm we can approximate the possibly incomplete successor candidate sets in a sufficient way. To ensure complete partitions we add a reference that covers all objects which cannot be referred to by the successor references (the *other* entry in our menu) as depicted in Eq. (3). Finally, the references can be reused in the selection process and therefore computed once.

The decision on which successor reference to use for refining the current selection is based on maximizing the resulting partition’s information content, which is similarly computed as in decision tree learning [42]. This strategy prefers to split the remaining objects in a way that reduces the total number of refinement strides. Moreover, the method allows to split the referable objects more equally, thus offering the user a meaningful choice at every stride. During the refinement process, we only offer choices that can result in an achievable goal, where goal reachability is efficiently approximated by *delete relaxation* [43]. For example, if all cups were out of reach of the robot, the choice *cup*(*x*) would be removed from the selection above. This can result in a completely different selection being preferred, e. g., one that uses the transportable’s color or position for distinction. If several objects satisfy the specified goal, the planner resolves this ambiguity by picking an arbitrary object among them.

### 4.3. Robot Motion Generation

For navigation planning of the mobile base, we apply the sampling-based planning framework *BI*^2^*RRT\** [44]. Given a pair of terminal configurations, it performs a bidirectional search using uniform sampling in the configuration space until an initial sub-optimal solution path is found. This path is subsequently refined for the remaining planning time, adopting an informed sampling strategy, which yields a higher rate of convergence towards the optimal solution. Execution of paths is implemented via a closed-loop joint trajectory tracking algorithm using robot localization feedback.

In this work, we additionally adopt a probabilistic roadmap planner approach [45] to realize pick, place, pour and drink motions efficiently. Therefore, we sample poses in the task space which contains all possible end-effector poses. The poses are then connected by edges based on a user-defined radius. We apply an A-based graph search to find an optimal path between two nodes using the Euclidean distance as the cost and heuristic function. To perform robotic motions we need to map the end-effector to joint paths, which can be executed by the robot. We thus use a task space motion controller which uses the robot’s Jacobian matrix to compute the joint velocities based on end-effector velocities. Additionally, collision checks ensures that there are no undesired contacts between the environment and the robot. Given a start- and end-pose of a manipulation task, the planner connects them to the existing graph and runs the mentioned search algorithm. For grasping objects, we randomly sample grasp poses around a given object and run the planner to determine a motion plan. Furthermore, we extract horizontal planes from the camera’s point cloud and sample poses on these planes to find a suitable drop location for an object. Finally, special motions as drinking and pouring are defined by specifying a path in the cup’s rim frame (the point which needs to be connected to the mouth during drinking) and the bottle’s rim frame, respectively. Based on these paths the planner samples roadmaps that allow to find motion paths close to the given ones in order to react to small changes in the environment.

### 4.4. Perception

This section outlines the perception techniques applied in this work and explains how objects and mouth locations are determined and liquid levels are estimated.

#### Object Detection

In order to detect objects we employ the method of Pauwels *et al*. [46] that relies on dense motion and depth cues and applies sparse keypoint features to extract and track six-degrees-of-freedom object poses in the environment. The algorithm additionally requires models that describe the structure and texture of the detectable objects. It is able to track multiple objects in realtime using a GPU-based solution. These poses and locations (e. g., the shelf) are finally stored and continuously updated in the knowledge base.

#### Pouring Liquids

An important aspect of pouring liquids, is to be able to determine when to stop pouring. This prevents overflowing and spilling the liquid as well as opening up possibilities such as mixing drinks, or preparing meals where exact amounts of liquid are required. Our approach to detect the liquid level employs an Asus Xtion Pro camera, which determines depth based on active structured light. Using this type of sensor, liquids can be categorized as either opaque or transparent. Opaque liquids, such as milk or orange juice, *reflect* the infrared light and the extracted liquid level represents the real liquid level (with some noise). In the case of transparent liquids, such as water and apple juice, the infrared light is *refracted* and the depth value is incorrect.

To detect the fill level of a transparent liquid, we base our approach on a feature further described by Hara *et al*. [47] and Do *et al*. [48]. This feature is given as follows:

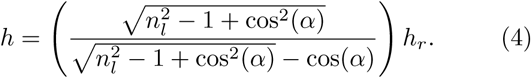

Here *h*_*r*_ represents the raw depth measured liquid level and *h* the estimated liquid height. The index of refraction of the liquid is given by *n*_*l*_ and angle *α* is the incidence angle of infrared light from the camera projector with respect to the normal of the liquid surface. A Kalman filter is then used to track the liquid level and compensate for noise.

Before pouring, we first detect the cup in the point cloud and determine a region within the cup boundaries where the liquid could be. During the pour, we extract the depth values for the liquid and estimate the real liquid height by either applying Eq. 4, in the case of transparent liquids, or using the extracted value directly, in the case of opaque liquids. The type of liquid and hence the index of refraction is given beforehand through the user’s selection. The viewing angle *α*, can be determined from the depth data. Once it is detected that the liquid level has exceeded a user defined value, a stop signal is sent to terminate the pouring motion.

#### Face Detection

We use a two-step approach to detect and localize the user’s mouth. In the first step, we segment the image based on the output of a face detection algorithm that uses Haar cascades [49, 50] in order to extract the image region containing the user’s mouth and eyes. Afterwards, we detect the position of the mouth of the user, considering only the obtained image patch. Regarding the mouth orientation, we additionally consider the position of the eyes in order to obtain a robust estimation of the face orientation, hence compensating for slightly changing angles of the head.

### 4.5. Dynamic Knowledge Base

The knowledge base provides data storage and establishes the communication between all components. In our work, it is initialized by a domain and problem description based on PDDL files. Once the knowledge base is initialized, it acts as a central database from which all participating network nodes can retrieve information about specific objects in the world as well as their attributes. Dynamic behavior is achieved by an additional layer that allows nodes to add, remove or update objects as well as their attributes. Moreover, the knowledge base actively spreads information about incoming changes as updates on object attributes across the network. Based on this information each network node decides on its own whether that information is relevant and which actions need to be taken.

## 5. Implementation Details

In our system, we distribute the computation across a network of seven computers that communicate among each other via ROS. The decoding of neuronal signals has four components. EEG measurements are performed using *Waveguard EEG* caps with 64 electrodes and a *NeurOne* amplifier in AC mode. Additionally, vertical and horizontal electrooculograms (EOGs), electromyograms (EMGs) of the four extremities, electrocardiogram (ECG), electrodermal activity (EDA) and respiration are recorded. The additional data is used to control for ocular and muscular artifacts, changes in heart beat frequency and skin conductance, and respiratory frequency, respectively. It is routinely recorded during EEG experiments in our lab. For recording and online-preprocessing, we use BCI2000 [38] and Matlab. We then transfer the data to a GPU server where our deep ConvNet, implemented using Lasagne [51] and Theano [52], classifies the data into five classes.

Furthermore, to find symbolic plans for the selected goal we use the A^*\**^-configuration of the *Fast Downward* planner [53]. The knowledge base is able to store objects with arbitrary attribute information. All changes in the knowledge base automatically trigger updates in the goal formulation GUI. Unexpected changes interrupt the current motion trajectory execution. Finally, we use *SimTrack* [46] for object pose detection and tracking and OpenCV for face detection.

## 6. Experiments

We evaluated the proposed framework in multiple experiments. Sec. 6.2 focuses on the BCI control of the whole system. Afterwards, the results regarding the goal formulation interface are presented in Sec. 6.3. We provide a detailed performance analysis (Sec. 6.3.2) and a user survey that studies the friendliness and intuitiveness of the goal formulation interface (Sec. 6.3.3) based on simulated environments with up to 100 objects in scenarios as exemplarily depicted in Fig. 9. Furthermore, we conducted two experiments in the real-world environment which are explained in Sec. 6.1. The results in Sec. 6.4.1 show that the framework is capable of handling fetch-and-carry tasks even if there are undesired changes in the environment. Finally, in Sec. 6.4.2 we discuss the results of combining all presented components into an autonomous robotic assistant that provides a drink to the user.

**Figure 9:**
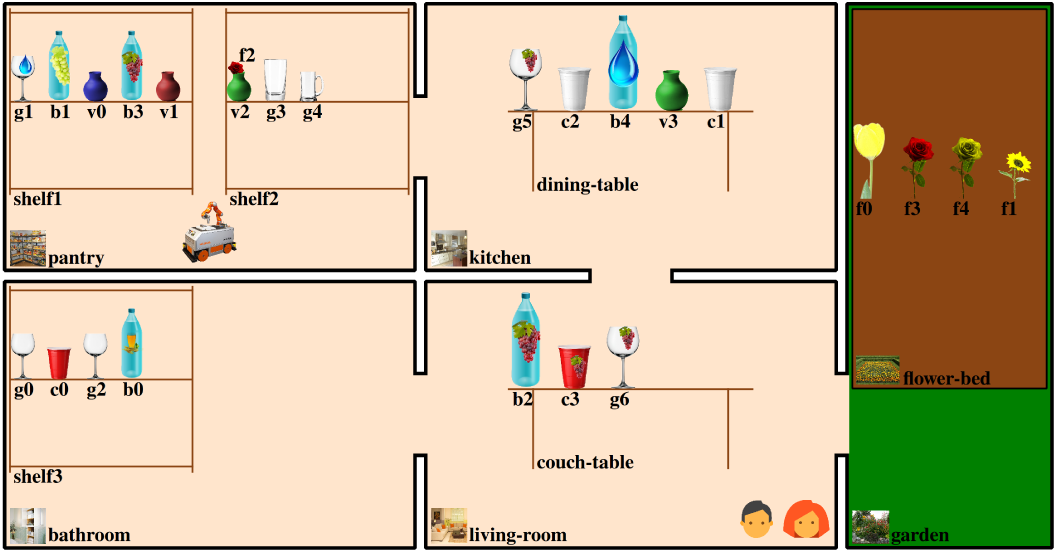
An exemplary scenario as used in our user experiments with four rooms, a garden, two humans, a robot and multiple objects.

### 6.1. Real World Experimental Environment

We performed multiple experiments in the real world scenario depicted in Fig. 7. It contains two shelves and a table as potential locations for manipulation actions. The user sits in a wheelchair in front of a screen that displays the goal formulation GUI. The autonomous service assistant we use is an *omniRob* omni-directional mobile manipulator platform by KUKA Robotics. The robot is composed of 10 DOF, i. e., three DOF for the mobile base and seven DOF for the manipulator. Additionally, a Schunk *Dexterous Hand 2.0* with three fingers is attached to the manipulator’s flange and used to perform grasping and manipulation actions, thus adding another DOF for opening and closing the hand. The tasks we consider in our experiments require the robotic system to autonomously perform the following actions: drive from one location to another, pick up an object, drop an object on a shelf or table, pour liquids from bottles into cups and supply a user with a drink. Moreover, our experimental setup uses a perception system composed of five RGBD cameras. Three of them are statically mounted at the shelves and the table, in order to observe the scene and to report captured information as object locations and liquid levels to the knowledge base. The other two cameras are carried by the robot on-board. The first one is located at the mobile base and used to perform collision checks in manipulation planning, whereas the second camera is mounted at the robot’s end-effector and used for tasks involving physical human-robot interaction as serving a drink to a user. Demonstrations of our work can be found online: http://www.informatik.uni-freiburg.de/~kuhnerd/neurobots/.

### 6.2. Online Decoding of Neuronal Signals

We evaluated the BCI control setup with four healthy participants (P1-4, all right-handed, three females, aged*±*26.75 5.9). In total, 133 runs have been recorded (90 with the real robot) where the participants selected various instructed goals and executed the corresponding task plans in the goal formulation GUI. For 43 runs, we used simulated feedback from the GUI in order to generate a larger amount of data for the evaluation. In this case, we simulated action executions by simply applying the corresponding effects to the knowledge base. Finally, 38 runs were discarded because of technical issues with the online decoding setup.

The performance of the BCI decoding during the remaining 95 runs was assessed using video recordings of interactions with the GUI. We rated GUI actions as correct if they correspond to the instructed path and incorrect otherwise. Actions which were necessary to remediate a previous error were interpreted as correct if the correction was intentionally clear. Finally, we rated *rest* actions as correct during the (simulated) robot executions and ignored them otherwise. For evaluation, five metrics have been extracted from the video recordings: (i) the accuracy of the control, (ii) the time it took the participants to define a high-level plan, (iii) the number of steps used to define a high-level plan, (iv) the path optimality, i. e., the ratio of the minimally possible number of steps to the number of steps used (e. g. 1 is a perfect path, while 2 indicates that the actual path was twice longer than the optimal path), and (v) the average time per step. We summarized the results in Table 1. In total, a 76.95 % correct BCI control was achieved, which required 9 s per step. Defining a plan using the GUI took on average 123 s and required the user to perform on average 13.53 steps in the GUI of the high-level planner. The path formed by these steps was on average 1.64 times longer than the optimal path, mainly because of decoding errors which had to be corrected by the participants, requiring additional steps. The decoding accuracy of every participant is significantly above chance (p *<* 10^−6^).

**Table 1:** Aggregated mean*±*std results for 95 BCI control runs (Exp. 6.2), p-value *<* 10

|  | Runs # | Accuracy [%]* | Time [s] | Steps # | Path Optimality | Time/Step [s] |
| --- | --- | --- | --- | --- | --- | --- |
| P1 | 18 | 84.1±6.1 | 125±84 | 12.9±7.7 | 1.6±0.6 | 9±2 |
| P2 | 14 | 76.8±14.1 | 90±32 | 10.1±2.8 | 1.1±0.2 | 9±3 |
| P3 | 28 | 78.8±9.5 | 173±144 | 16.9±11.5 | 2.1±1.3 | 10±4 |
| P4 | 35 | 68.1±16.3 | 103±69 | 14±7.6 | 1.7±0.74 | 7±2 |
| | 95 | 76.9±9.1 | 123±36 | 13.5±2.8 | 1.6±0.4 | 9±1 |

The participant-averaged EEG data used to train the hybrid ConvNets and the decoding results of the train/test transfer are visualized in Fig. 8. In Fig. 8(a) we show the signal-to-noise ratio (SNR) of all five classes of the labeled datasets. We define the SNR for a given frequency *f*, time *t* and channel *c* as

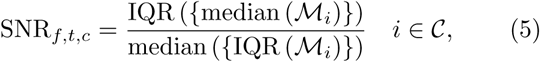

where *ℳ*_*i*_ corresponds to the set of values at position (*f, t, c*) of the *i*-th task, with |ℳ_*i*_ *|* being the number of repetitions. median(·) and IQR(·) is the median and interquartile range (IQR), respectively. The upper part describes the variance of the class medians, i. e., a large variance means more distinguishable class clusters and a higher SNR. The denominator corresponds to the variance of values in each class, i. e., a lower variance of values results in a higher SNR.

In all non-peripheral EEG electrodes a clear and sustained increase in SNR is visible in the alpha (*∼*8-14 Hz) and beta (*∼*14-30 Hz) frequency bands, starting around 500 ms after the cue. These frequency bands are robust markers of brain activity. The partial absence of the increased beta band SNR in peripheral channels further supports the neuronal origin of the signal [54]. An increased SNR is also visible in both EOG channels which could indicate a contamination of the EEG data by ocular artifacts. The slight increase in SNR in the horizontal EOG channel in the delta (*∼*0-4 Hz) and theta (*∼*4-8 Hz) bands 0.5-1 s after the cue is most probably due to residual *neuronal* activity recorded by the EOG. Support for this assumption is based on the fact that this increase is stronger in most EEG electrodes, suggesting a generator located some distance from the eyes, i. e., in the brain. The sustained increase in SNR in the delta band visible in the vertical EOG is likely due to unconscious eye movements. As this increase in the delta band SNR is only visible in the three front-most EEG electrodes and weaker than the increased SNR of unambiguous neuronal origin described above, we are confident that the hybrid ConvNets will not have learned to use this activity to differentiate between the mental tasks. The visualizations shown in Fig. 8(c) support this idea as no correlations are visible for frontal EEG electrodes in the delta band. The increased SNR in the lower frequencies of the respiration and EDA channels is probably related to task engagement. A crosstalk between these signals and the EEG is unlikely and not supported by the SNR analysis. The extremely low SNR in all EMG channels shows that the participants performed pure imagery, without activating their limb muscles. In summary, the SNR analysis revealed that the offline training data contains informative neuronal activity which the hybrid ConvNets should have been able to learn from.

Indeed, the decoding accuracies (mean 63.0 %, P1 70.7 % increasing the power in the given frequency bands at the P2 49.2 %, P3 73.1 %, P4 58.8 %) resulting from the test dataset after initial training of the ConvNets are well above the theoretical chance level of 20 %. These are visualized in Fig. 8(b) in the form of a pooled confusion matrix. Right hand and feet motor imagery were most often confused with each other, mental rotation was evenly confused with all other classes and word generation and rest were most often confused with feet motor imagery. The co-adaptive online training which took place between the initial training of the ConvNets and the online evaluation increased the decoding accuracy from 63.0 % to 76.9 %, which is a clear indication for the efficacy of our approach. It should further be noted that the increase in accuracy occurred from an *offline, cued* evaluation to an *online, uncued* evaluation, which is quite remarkable. It has to be mentioned however that the online accuracy is a subjective measure as the intentions of the participants had to be inferred from the instructions (cf. Sec. 4.1.2). The offline accuracy was fully objective because of the presented cues. Nevertheless, the online evaluation decoding accuracy leaves room for improvements. Preliminary offline steps have been undertaken using the data collected during the offline and online co-adaptive training to detect decoding errors directly from the neuronal signals [20]. This first attempt already yielded mean error detections of 69.33 %. The detection accuracy could potentially be increased by including error sensitive peripheral measures as EDA, respiration and ECG into the decoding. Access to the high-gamma (*∼*60-90 Hz) band frequency range could further increase the decoding accuracy of both mental tasks [15] and error signals [55]. Once transferred to an online experiment one could use this error detection to undo the error, generate a new decoding and retrain the decoder. Lastly, detection of robotic errors could also be achieved from the ongoing EEG [56, 57, 58, 21] and used as both emergency stop and teaching signals.

To further support the neural origin of the BCI control signals, Fig. 8(c) shows physiologically plausible input-perturbation network-prediction correlation results (see [15] for methods). Specifically, predictions for right hand and feet motor imagery classes were negatively correlated with input-perturbarions (see [15]) in the alpha and beta bands at EEG electrodes located directly above their motor and somatosensory cortex representations. This means that specific electrodes resulted in reduced predictions. By symmetry, reduced power resulted into increased predictions. These correlations fit well with the neuronal basis of the event related desynchronisation of the alpha and beta bands during motor imagery [59]. A positive correlation is also apparent above the foot motor and somatosensory cortex for feet motor imagery in the delta band. This positive correlation probably reflects the feet motor potential [60]. For mental rotation, word generation and rest the input-perturbation network-prediction correlation results are less easily interpretable, mostly due to the lack of extensive electrophysiological reports. A positive correlation is visible for the mental rotation above the medial parietal cortex in the alpha band which could reflect the involvement of cortical representations of space. Similarly, positive correlations are visible bilaterally above the lateral central cortex and temporal cortex in the low beta band during word generation. They could reflect the involvement of speech and auditory brain areas. Further investigations will be needed to delineate these effects.

### 6.3. Goal Formulation Interface

In this section we present a performance experiment to evaluate the runtime required by the GUI and the results of a preliminary user study, which examines the user-friendliness and intuitiveness of the system. Moreover, we discuss how humans use references to objects.

#### 6.3.1 Scenario Setup

We created a virtual scenario with five rooms as depicted in Fig. 9: a kitchen with a dining table, a living room with a couch table, a pantry with two shelves, a bathroom with one shelf and a garden containing a flowerbed. Bottles, cups, glasses and vases are distributed among the furniture. There are three types of flowers (e.g., *rose*), seven drinking contents (e.g., *red* -*wine*), five colors (e.g., *red*) for cups and vases and three for flowers and finally, four glass shapes (e.g., *balloon*). Flowers can be put into vases but may also be placed directly on furniture. The *omniRob* robot has the ability to move between the rooms and serve the two persons (*me* and *friend*). Finally, the available actions are: *arrange* a flower in a vase, *pick* a flower out of a vase, *grasp* and *drop* an object, *give* an object to a human, *pour* a liquid from one vessel to another, *drink* to assist a human with drinking a drink, *move* the robot between rooms and *approach* a furniture or human for further interaction.

#### 6.3.2 Performance

In this experiment we evaluated the performance of the goal formulation interface. We used a scenario generator which randomly creates instances of the planning problem. To assess the performance, we measured the time required to start the user interface and select parameters of random actions. The experiment was repeated 100 times and averaged to retrieve reliable results. The performance of our Python implementation was determined using an Intel i7-3770K (3.5 GHz) processor and 16 GB of memory. Fig. 10 illustrates the run times needed for several operations as a function of the number of objects present in the world. The most time-consuming component is given by the reference exploration, where initial partitions are chosen (Fig. 10 left, red). Another computationally expensive operation is the root menu generation, which determines potentially reachable goals for all actions based on delete relaxation (Fig. 10 left, green). In contrast, the reference refinements for the current parameter of an action requires in average less than 2 s even for scenarios containing numerous objects (Fig. 10 right). However, this assertion only holds as long as the world and thus the references do not change. Considering dynamic environments, changes of the world are frequently triggered by, e. g., actions taken by the robotic service assistant. For example, when the robot has grasped a cup, the system should no longer refer to the cup as *the cup on the table*. Instead, the reference must be rebuilt given the updated environment state yielding *the cup at the robots gripper*. For simplicity, our approach rebuilds all object references when an environment change has been detected. In the future, only obsolete references should be recomputed in order to scale well on larger scenarios.

**Figure 10:**
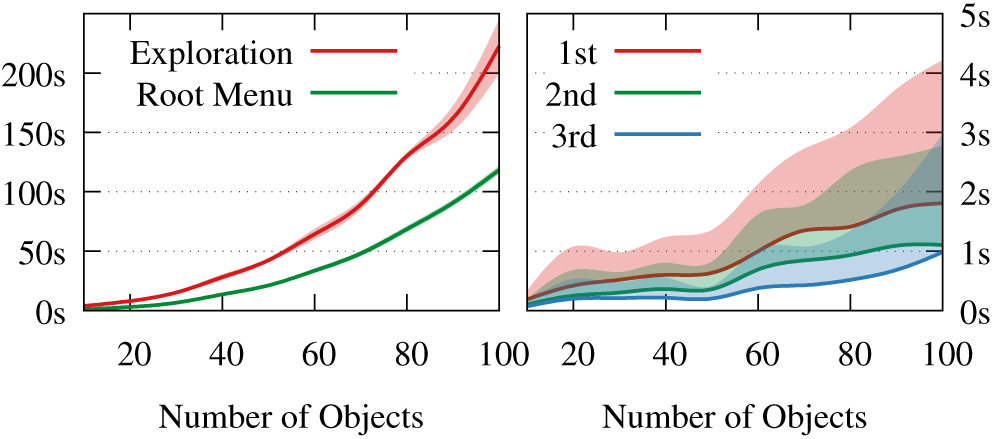
Evaluation of the computation time for different numbers of objects in the environment averaged over random actions. *Left:* The plot shows the mean and standard deviation of building the menu structure at the beginning and includes initial exploration and root menu creation. *Right:* Refinements of a goal can be done efficiently. It shows the mean and positive standard deviation times of the first three refinements.

#### 6.3.3. User Study

##### Participants

A total of 20 participants (3 female, 17 male, 25 – 45 years) took part in the user study and gave their consent for the anonymized processing of the collected data. The participants were students in computer science and administrative employees of the university. They used our system the first time and were not familiar with it.

##### Data Collection and Measures

The participants had to use our system to accomplish tasks in five simulated scenarios, which were generated beforehand to get comparable results. The five scenarios with increasing complexity were: (S1) Move the robot to the garden, (S2) Drink beer using a beer mug, (S3) Arrange a red flower in a red vase, (S4) Place a red rose on the couch table, and (S5) Give a red wine glass with red wine to your friend. After introducing the user interface by explaining the individual components of the system, the participants had to accomplish the five tasks using the GUI. Since there were no time constraints and sub-optimal strategies were allowed, all users managed to reach the requested goal states. We1 counted the number of *steps* the participants required to finish the predefined tasks successfully, where a step is either a refinement of an attribute or the selection of the *back* entry in the menu.

For each scenario the participants had to rate if the displayed control opportunities offered by the user interface *comply* to their expectations in a questionnaire, where the *compliance* levels ranged from 1 (unreasonable) to 5 (fully comply). Moreover, we asked the participants to rate the overall *intuitiveness* of the GUI in the range of 1 (not intuitive) to 5 (excellent). We then asked whether the participants prefer to describe objects using references or via internal names (e.g., *v2*). Additionally, we evaluated the *subjective quality* of object references ranging from 1 (not prefer at all) to 5 (highly prefer). We proposed four references to objects depicted in Fig. 9 and let the users rate how well each of those references describes the corresponding object. Moreover, subjects were asked to generate references to these objects in natural language themselves in the way they would tell a friend to find an object. In particular, we considered the green vase with the red rose located in the pantry (*v2*) and the glass, filled with red wine (*g6*), located on the couch table in the living room. The proposed references ranged from under-determined to over-determined descriptions, e.g., *the green vase* vs. *the green vase located in the right shelf in the pantry which*1 *contains a red rose*.

*Result* Fig. 11 shows the quantitative result of the user study. We counted the number of steps performed by each of the participants to achieve the predefined tasks successfully. The figure shows box plots for each scenario. Additionally, the plot contains the optimal number of steps which are required to successfully achieve the goal. Most of the participants were able to find a near-optimal strategy to solve the task. The outliers in the first four scenarios are mainly caused by the user exploring the possibilities of the user interface. The increased number of steps in the last scenario can be traced back to the following reasons. First, the scenario required two actions to be able to achieve the task: fill a balloon shaped glass with red wine and give this glass to the friend. Only a few users were able to determine this fact at the beginning. Therefore, the participants had to correct their decisions which results in a higher number of steps in the fifth scenario. Second, the pour action as defined in our scenarios required to specify three parameters: the vessel to pour from, the vessel to pour to and the liquid that is poured. Our system usually refers to the first vessel by its content, so the redundant refinement of the liquid as last parameter is not intuitive to the users. Finally, we split a partition based on its information content to reduce the number of refinements. This strategy can lead to unexpected refinements of object attributes since the user might prefer these in a different order.

**Figure 11:**
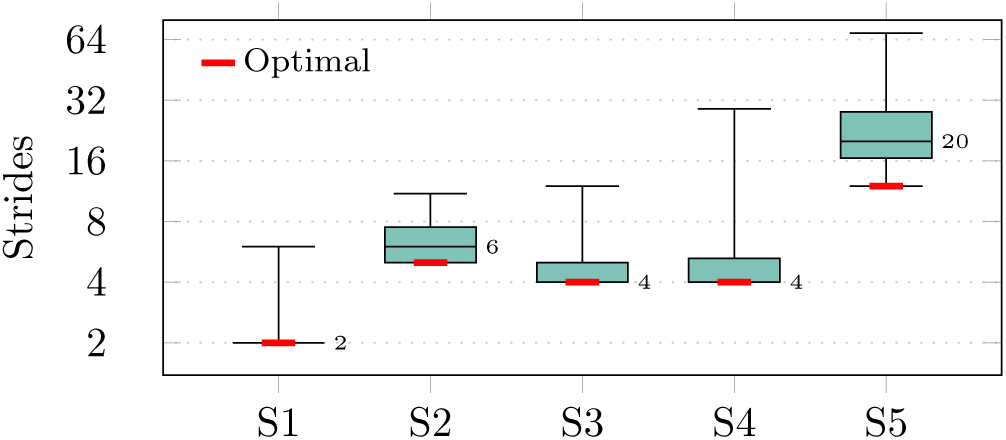
The box plots illustrate the number of steps required by our participants to achieve a given goal in five different scenarios S1-S5 (optimal number of steps indicated in red, numbers denote the median)

Fig. 12 shows the results on how well the choices offered by the high-level planning GUI actually comply with the expectations of the users. A large percentage of them comply with the refinements provided by the GUI in the scenarios S1 to S4. Due to the previously mentioned problems however, S5 has been rated worse. A short training period of the users to get familiar with the interface might help to improve the compliance in S5. Overall, 80% of the participants rated the GUI as intuitive, i.e., according to the aforementioned metric they rated the intuitiveness with at least 3 (acceptable). In particular, 85% of the participants preferred referring to objects by incremental referencing over internal names (e.g., *green vase on the couch table* vs. *v1*).

**Figure 12:**
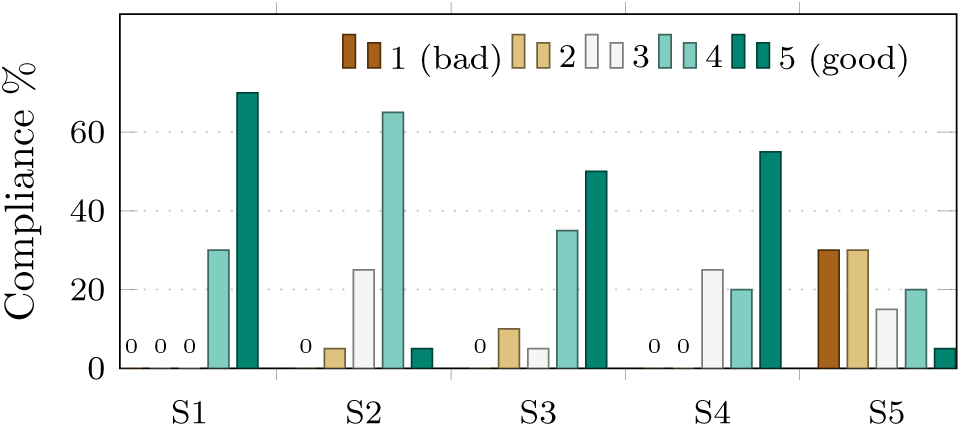
Compliance of the offered choices with the users’ expectation for five tasks in different scenarios. The participants had to1 select compliance levels from 1 (unreasonable) to 5 (fully comply).

In the last user experiment, we evaluated the *subjective quality* of object references. According to our results, preferred references highly depend on whether the spatial context of the agents in the world is considered or not. One group of users only preferred references that uniquely identify the objects independent from the location of the agents. This group preferred references such as *the vase containing a rose* or occasionally also *the vase in the right shelf* for *v2* and *the red wine glass on the couch table* for *v6*. Another group preferred under-determined references as they considered the spatial context of the agents. This group preferred references such as *the green vase* for *v2* and *the red wine glass* for *v6*. Interestingly, the capability of users to impersonate the acting agent has also a strong influence on the references preferred by the second group. For referring to *v2*, some users of the second group additionally specified the room or the content of the vase, assuming that the assisting agent is also located in the living room and therefore requires a more detailed object description, while they preferred under-specified reference for objects on the couch table. Detailed over-specified references were refused by all participants, but more firmly b the second group. Summarizing, our evaluation revealed that incrementally building object references is suitable to describe objects precisely. Room for improvement was identified in updating object references that change during plan execution and in the consideration of temporal and spatial context.

### 6.4. Robotic Service Assistant

We performed two experiments to evaluate the system in the real world using a mobile robot. The first one explores how the system reacts to unexpected changes. In the second experiment, we present the results of the whole system involving all components.

#### 6.4.1 Fetch and Carry Task with Disturbances

In a dynamic world, unexpected changes such as adding or removing objects can occur at all times. With this experiment, we examine how our system adapts to disturbances, i. e. unexpected changes in the environment.

We performed the experiments in a way that unexpected world changes may occur at any time through actions taken by another unknown agent. In practice, this agent could refer to a human taking actions that directly affect the execution of the current high-level plan. Therefore, we initially placed a cup on one of the shelves and queried the goal formulation assistant to generate a sequence of actions leading to the goal state *cup on table*,. i. e., *approach*(*shelf with cup*), *grasp*(*cup*), *drop*(*cup*). *approach*(*table*), Once the robot arrived at the corresponding shelf in the execution phase of the plan, a human agent took the cup while the robot was about to grasp it and transferred it to the other shelf. In order to obtain quantitative results on the performance of our framework in such a scenario, we ran this experiment 10 times with different initial cup placements and evaluated its ability to generate the goal state in the real world despite the external disturbance introduced by the human agent. For all runs, our perception system correctly updated the information on the cup in the knowledge base, in turn triggering a re-planning step. The updated action sequence always contained two additional actions, namely moving and standard deviation is 6.9 8.9 mm. In some instances the bottle obstructed the camera view, resulting in poor liquid level detection and a higher error. We have begun investigation possible improvements for the monitoring of liquid levels by additionally considering the brain activity of an observer [58, 21]. Our latest results show that events where the liquid spills over the cup can be detected with an accuracy of 78.2*±*8.4 % (mean over 5 subjects) using to the shelf where the human agent dropped the cup and grasping it again. In total, 59 out of 60 (98.33%) scheduled actions were successfully executed and thus 90% of the runs succeeded in generating the goal state. Only one run failed in the action execution phase due to the inability of the low-level motion planning algorithm to generate a solution path for the mobile base within the prescribed planning time. On average, our system required an overall time of 258.7*±*28.21 s for achieving the goal state.

#### 6.4.2 Drinking Task

The last experiment evaluates the direct interaction between user and robot. Therefore, we implemented an autonomous robotic drinking assistant. Our approach enabled the robot to fill a cup with a liquid, move the robot to the user and finally provide the drink to the user by execution of the corresponding drinking motion in front of the user’s mouth. Fig. 13 shows examples for the actions *move, grasp* and *pour*. The first row contains the task-plan visualizations of the goal formulation GUI, which are displayed after a goal has been selected. Additionally, the second row depicts the planning environment as used by the navigation and manipulation planners to generate collision-free motions. The corresponding view of the real world is shown in the last line.

**Figure 13:**
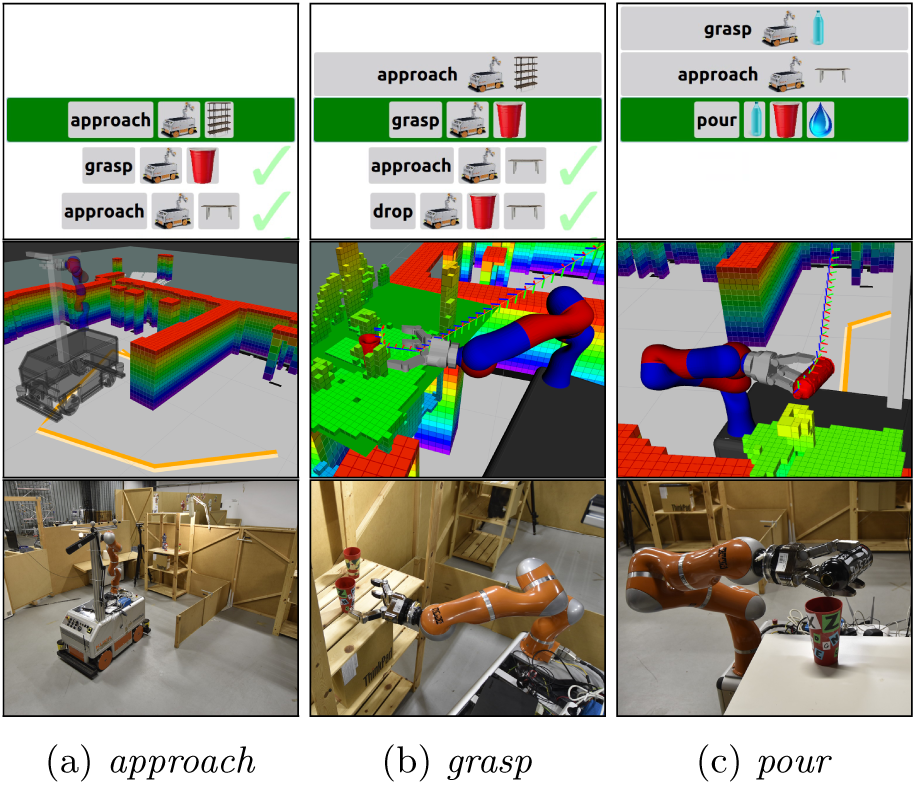
Snapshots of our experiments for the actions *approach, grasp* and *pour*. The first line shows the corresponding step in the high-level planner user interface. The results of the motion and manipulation planning is depicted in the second row. Finally, the third row shows the robot system, which executes the actions.

Table 2 shows the averaged results for the experiment. Again, the user is one of the authors. Here, only 3.75 % of the 160 scheduled actions had to be repeated in order to complete the task successfully. In one run, plan recovery was not possible leading to abortion of the task. Thus, our system achieved in total a success rate of 90 % for the drinking task. Planning and execution required on average 545.56 67.38 s. For the evaluation of the liquid level detection approach, we specified a desired fill level and executed 10 runs of the pour action. The resulting mean error deep ConvNets and EEG [21]. This feedback can be used to inform the liquid level detection and pouring procedure of the failure and trigger an adaptation of the algorithms to prevent future spills. To completely prevent errors, detection prior to a spill event will have to be achieved in future work.

**Table 2:**
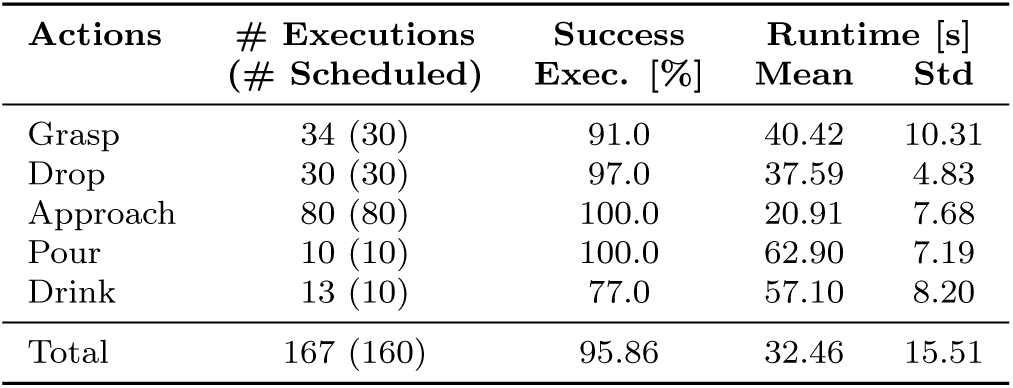
Aggregated results for 10 runs (Exp. 6.4.2)

| Actions | # Executions<br>(# Scheduled) | Success<br>Exec. [%] | Runtime [s] |  |
| --- | --- | --- | --- | --- |
|  |  |  | Mean | Std |
| Grasp | 34 (30) | 91.0 | 40.42 | 10.31 |
| Drop | 30 (30) | 97.0 | 37.59 | 4.83 |
| Approach | 80 (80) | 100.0 | 20.91 | 7.68 |
| Pour | 10 (10) | 100.0 | 62.90 | 7.19 |
| Drink | 13 (10) | 77.0 | 57.10 | 8.20 |
| Total | 167 (160) | 95.86 | 32.46 | 15.51 |

## 7. Conclusions

In this paper, we presented a novel framework that allows users to control a mobile robotic service assistant by thought. This is particularly interesting for severely paralyzed patients who constantly rely on human caretakers as some independence is thereby restored. Our system performs complex tasks in dynamic real-world environments, including fetch-and-carry tasks and close-range human-robot interactions. Our experiments revealed that the five-class-BCI has an uncued online decoding accuracy of 76.9 %, which enables users to specify robotic tasks using intelligent goal formulation. Furthermore, a user study substantiates that participants perceive the goal formulation interface as user-friendly and intuitive. Finally, we conducted experiments in which the proposed autonomous robotic service assistant successfully provides drinks to humans. By combining techniques from brain signal decoding, natural language generation, task planning, robot motion generation, and computer vision we overcome the curse of dimensionality typically encountered in robotic BCI control schemes. This opens up new perspectives for human-robot interaction scenarios.

## 8. Acknowledgements

This research was supported by the German Research Foundation (DFG, grant number EXC1086), the Federal Ministry of Education and Research (BMBF, grant number Motor-BIC 13GW0053D) and grant BMI-Bot (grant number ROB-7) by the Baden-Württemberg Stiftung.

## Footnotes

2 We initially used 1 s intervals to maximize speed, but they were too short for proper mental task transition.

3 ^3^Initially we defined significance as *p <* 0.1. Initial experiments however showed that the time required for accumulating evidence to push *p* from 0.2 to 0.1 was disproportionally large. We therefore define significance as *p <* 0.2 to speed-up the decoding at the cost of accuracy.

